# Measurement of plant water status via static uniaxial compression of the leaf lamina

**DOI:** 10.1101/2022.03.29.486324

**Authors:** Tomás I. Fuenzalida, Oliver Binks, Callum J. Bryant, Joe Wolfe, Marilyn C. Ball

## Abstract

Turgor pressure is an essential, but difficult to measure indicator of plant water status. Turgor has been quantified by localised compression of cells or tissues, but a simple method to perform these measurements is lacking. We hypothesized that changes in leaf turgor pressure can be monitored by uniaxially compressing the leaf lamina and measuring the mechanical stress under a constrained thickness (stress relaxation); and that changes in leaf water content can be monitored by measuring the thickness of the leaf lamina compressed under a constant force (creep). Using a custom-built leaf squeeze-flow rheometer, we performed different compression tests on leaves from thirteen plant species. The equilibrium mechanical stress measured during stress relaxation was correlated with leaf turgor pressure (R^2^ > 0.95) and thus with leaf water potential (R^2^ > 0.94); the equilibrium leaf thickness measured during creep was correlated with relative water content (R^2^ > 0.74). The coefficients of these relationships were related to the leaf osmotic pressure at the turgor-loss point. An idealised average-cell model suggests that, under isothermal conditions, the bulk cell stiffness during compression is largely determined by the leaf osmotic pressure. Our study presents an inexpensive, accessible and automatable method to monitor plant water status non-invasively.

## Introduction

Plants live in pulses. The oscillations that occur in the physical environment cause large variations in growth rate and photosynthetic activity. Such variations are often related to changes in the hydrostatic pressure of cells, called turgor pressure (P). Turgor gives some rigidity and structure to soft plant tissues, such as leaves, flowers and growing stems and roots, and causes the swelling and shrinking movements that occur with changing water content (Skotheim Jan and Mahadevan, 2005). Aside from its structural role, turgor supports growth by driving plastic expansion of the cell wall (Lockhart, 1965); affects organ development through complex mechanical feedback loops (Green, 1962, Hamant et al., 2008); and causes stomata to open (Franks et al., 1998), regulating the rates of gas exchange between leaves and the atmosphere. Turgor is thus essential to plant life, and is also a valuable indicator of plant water status. However, turgor measurements are not routine.

Three main approaches to measure turgor pressure exist. First, turgor can be measured directly, by probing and measuring the hydrostatic pressure inside a single cell (Green and Stanton, 1967, Hüsken et al., 1978). This technique also enables the quantification of other cell parameters, such as the bulk elastic modulus and the membrane hydraulic conductivity. However, the method is relatively complex and requires careful manipulation, making it largely impractical in field settings. In a second approach, turgor can be inferred from the relation between water content and water potential (Ψ) of plant tissues (Tyree and Hammel, 1972). When leaves are wilted (*i*.*e*., turgor is nearly zero), the balancing pressure (BP = –Ψ) required to push water out of a cut leaf increases in proportion to the intracellular concentration of solute (Scholander et al., 1965). By measuring the leaf balancing pressure and water volume during slow dehydration, an estimate of the osmotic pressure^1^ (Π) at all levels of hydration can be obtained, whereby the turgor pressure can be derived as P = Ψ + Π. This widely used technique is simple and accurate, but cannot be performed on tissue still attached to the plant, is time consuming and laborious, and cannot be easily automated.

A third approach to measure turgor pressure is via indentation or compression of cells or tissues. Various techniques can be employed for this purpose; they have been reviewed by Geitmann (2006), Beauzamy et al. (2014) and Bidhendi and Geitmann (2019). The main principle behind these measurements is that the mechanical properties can be derived and used to infer hydration status by measuring the force and the deformation applied. While simple in principle, these techniques are commonly applied at the nm-µm scale and require costly instrumentation such as micro-indenters or atomic-force microscopes. Additionally, they often need to employ optical microscopy to determine the contact area between the sample and the indenter, which is necessary to obtain the stress. Similar principles could be applied to tissues at a larger (µm-mm) scale, where the cost of instrumentation can be reduced significantly. Such measurements could be useful for environmental plant physiologists and agronomists, who will mostly be interested in bulk tissue properties. Thus, the goal of this study is to test if these principles can be scaled up to produce a more practical and affordable method to monitor plant water status.

### Leaf squeeze-flow rheometry

There are various names given to techniques that compress samples between parallel plates, such as ‘simple unconfined compression’, ‘uniaxial compression’ and ‘squeeze-flow’ (Engmann et al., 2005). The disciplines which have coined these terms fall broadly within the fields of material science and rheology. Rheology is the science that studies the flow of matter under load, with particular emphasis on viscoelastic and plastic materials. Rheological responses usually involve time-dependent deformation due to their viscous or plastic components. At the tissue scale, the rheology of fruits and vegetables has been linked to turgor pressure (Lin and Pitt, 1986, De Belie et al., 2000, Jackman et al., 1992); however, this has not been applied to monitor plant water status. Some measurement paradigms from rheology could be useful to plant scientists studying plant hydration dynamics, so we refer to the technique of uniaxially compressing leaves as ‘leaf squeeze-flow rheometry’. Although transport phenomena relevant to squeezing leaves involve bulk flow and also diffusion, the term is adopted here in the spirit of encouraging interdisciplinarity between plant science and rheology. Here we introduce a first approach that may inspire further development by rheologists and plant scientists.

Two standard experimental paradigms in rheology are creep and stress relaxation. In a creep experiment, a sample is loaded with a constant stress (the force per unit area), and the resulting deformation is monitored over time; in a stress relaxation experiment, the sample is constrained to a given deformation, and the change in stress is monitored over time. We hypothesized that creep and stress relaxation could be used as measurement paradigms to monitor leaf water content and leaf turgor pressure, respectively. (The concepts here are somewhat different from standard rheology, because the mass of the leaf tissue studied is usually not conserved.) Our working premises were simple: to monitor changes in turgor pressure, the normal stress applied to the compressed leaf may be measured under a constrained leaf thickness; to monitor leaf water content, the thickness of the leaf may be measured under a constant load. An illustration of our working hypotheses is shown in Fig. 1 for two different measurements during a change in leaf hydration. (In this paper, the imposed mechanical stress is always compressive. By convention, tensile stresses are positive, so the applied compressive stress −σ is used throughout.)

**Figure 1.**
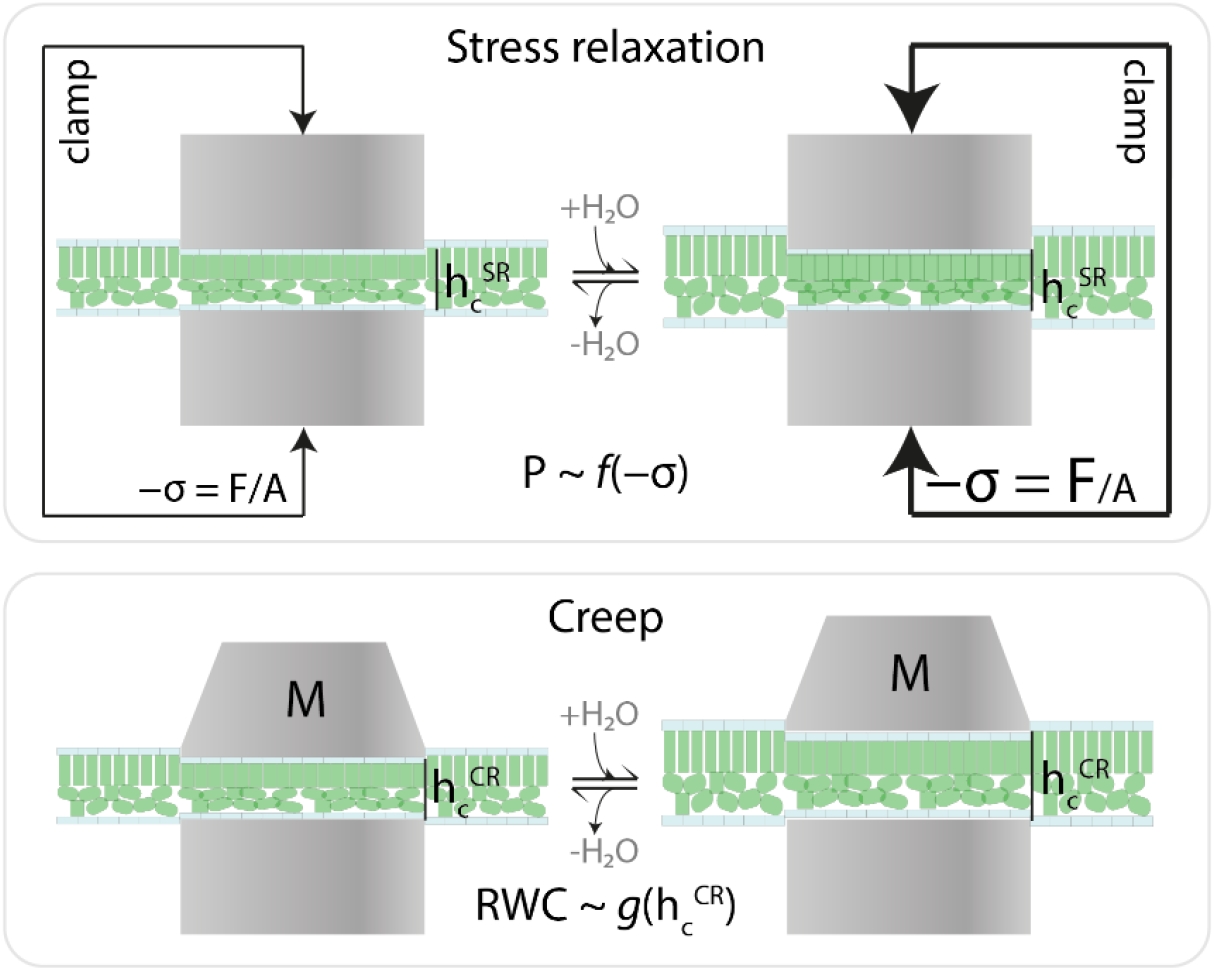
Illustration of our working hypotheses. In a stress relaxation experiment (top row), the leaf is compressed with a clamp to a constrained leaf thickness (h_c_^SR^); as the leaf changes water status (depicted by the rehydration/dehydration arrows), we hypothesise that leaf turgor pressure (P) can be approximated as a function *f* of the applied compressive stress (−σ = F/A). (The size of the arrows represents the magnitude of the clamping force under changing water status, which under constrained thickness changes due to the reaction force that the leaf exerts on the clamp.) In a creep experiment (bottom row), the leaf is compressed between two parallel plates under constant force, here represented by a weight of mass M; as the leaf changes water status, we hypothesise that leaf relative water content (RWC) can be approximated as a function *g* of the compressed leaf thickness h_c_^CR^.

## Methods

### Leaf squeeze-flow rheometer

Experiments were conducted using an affordable (∼US$300) custom-made leaf squeeze-flow rheometer (Fig. 2). Variations on the design of the instrument occurred during this study; in this is section we describe the version shown in Fig. 2 (differences are mentioned where appropriate). A load cell (FX29, TE Connectivity, Switzerland) was mounted between two microscope slides held together by two rubber bands and placed between the anvil and spindle of a 500 µm pitch analog micrometer (103-129, Mitutoyo, Japan). Leaf samples were loaded between the micrometer anvil (diameter = 6.4 mm) and the upper microscope slide to minimise shear stress caused by the spindle rotation. A microcontroller (Arduino MEGA 2560, Keyestudio, China) performed pressure readings from the load cell and operated a two-phase 400 steps/revolution NEMA17 stepper motor using a stepper motor driver (TB6600, China). The motor was coupled to the micrometer knob using a rigid shaft. Because the micrometer knob translates with the spindle rotation, the motor was mounted on a sliding linear guide to allow motor translation. The motor was operated at a microstep setting of 1/32 of a step, giving a theoretical minimum translation step of *c*. 39 nm. Translation of the micrometer spindle was calculated by counting number of steps taken by the motor. A digital temperature sensor (BME280, Core Electronics, Australia) was directly and firmly attached to the micrometer handle and insulated using a closed-cell polyurethane foam to monitor the temperature of the frame. An infrared digital temperature sensor (SEN0206, DF Robot, China) was used to track the leaf temperature.

**Figure 2.**
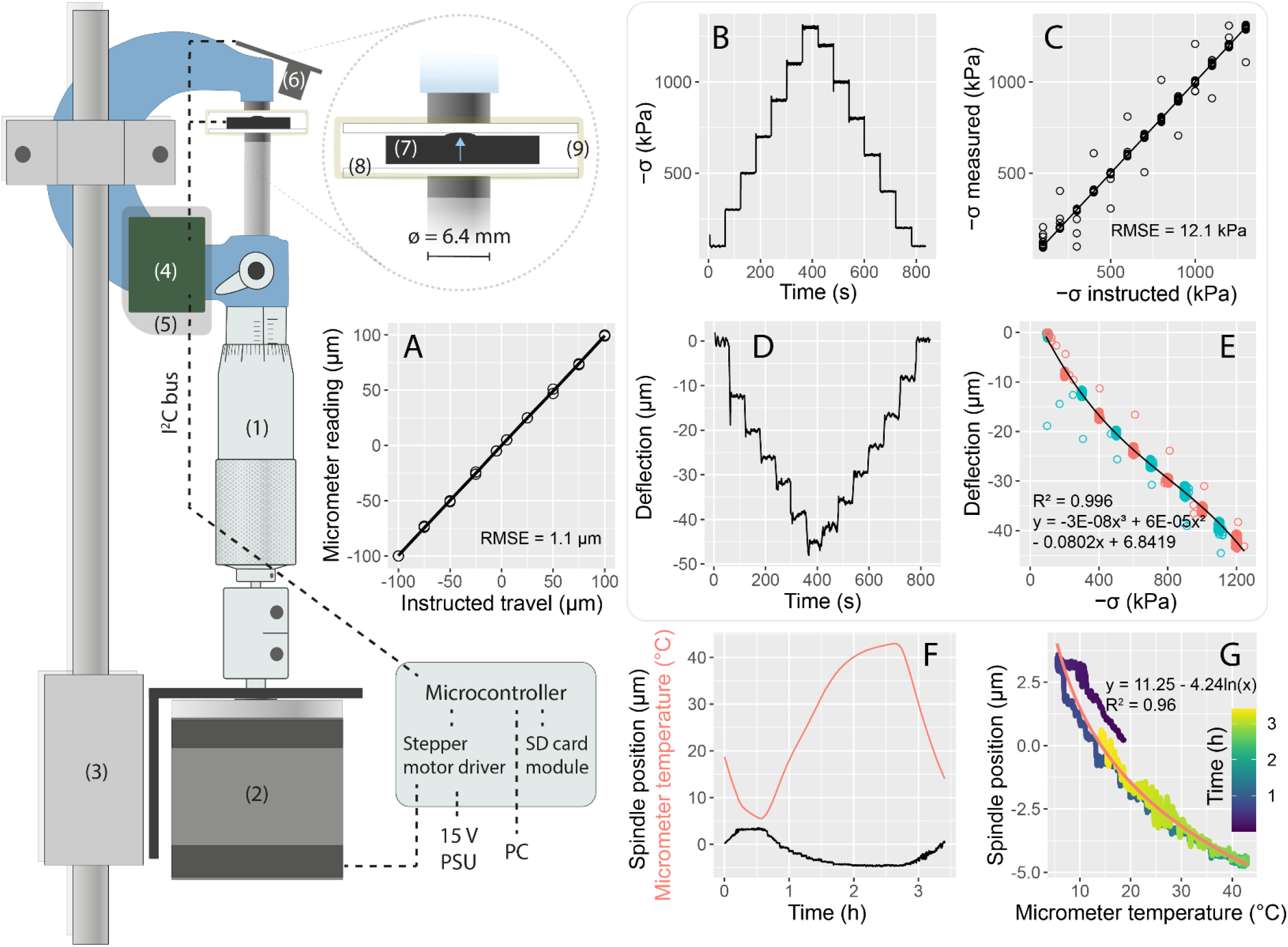
Component diagram and calibration of the leaf squeeze-flow rheometer. Components: (1) analog micrometer; (2) NEMA17 stepper motor; (3) linear guide; (4) micrometer temperature sensor; (5) foam insulation pad; (6) infrared leaf and ambient temperature sensor; (7) button compression load cell; (8) microscope glass slide; (9) rubber band. Leaf samples were compressed between the upper microscope slide and the micrometer anvil. Force, displacement and temperature data were logged to a SD card. Dashed lines denote simplified connections between components. A: Relation between the instructed distance to travel and the micrometer reading. B: Compressive stress applied (−σ = F/A) during a stairs creep test on an empty system (compressing the upper microscope slide against the micrometer anvil) with 200 kPa steps. C: Relation between the stress instructed and the stress measured by the load cell during the stairs creep test shown in B. D: System deflection during the creep test depicted in B. E: System deflection under load during the creep test shown in B; red and blue symbols denote measurements during increasing and decreasing load. F: Curves illustrating temperature sensitivity of the empty instrument during a thermal cycle while performing a creep test at 100 kPa: red is the temperature imposed and black the displacement recorded simultaneously. G: Relation between the micrometer temperature and the spindle position during the thermal cycle shown in F.

The instrument was programmed to perform creep and stress relaxation (SR) experiments. For creep experiments, the motor was instructed to move in either direction in response to the pressure reading difference from the set point (*i*.*e*., negative feedback). The motor step size was varied according to the predicted mechanical compliance of the empty system (Fig. 2E) as a function of the instantaneous error between the set point and the process variable. This control system can be described as a form of proportional ‘bang-bang’ control.

For the stress relaxation experiments, the motor was programmed to move in proportion to the estimated deflection of the load cell, or to not move at all. We distinguish these tests as SR ‘*sensu stricto*’, and SR ‘*sensu lato*’, respectively. As compression decreases during SR, the system dimensions change, so the micrometer handle must be turned to compensate the deflection and maintain the sample thickness close to constant (ASTM E328-13). Briefly, this was achieved by calculating the predicted mechanical compliance (Fig. 2E) as a function of the change in pressure between consecutive readings during SR, and by correcting this deflection by taking discrete steps. As positioning corrections were always a multiple of the minimum step size (*c*. 39 nm), corrections introduced an error given by the remainder between the step size and the estimated system deflection. This positioning error was added over time and corrected in the next control cycle, providing effective control over the sample thickness during SR.

The type of measurement just described, stress relaxation *sensu stricto* (SR_SS_), is desirable because it enables measurements to be reproducible independent of the instrument’s mechanical compliance, although it can be difficult to achieve and may introduce noise to the data. The present instrument can only effectively perform SR_SS_ tests when the direction of stress relaxation does not change. Changing direction involves taking ‘dead steps’ due to the instrument backlash, and this issue is not corrected for in the present version of the instrument; in most cases, positioning corrections during SR_SS_ are much smaller than the instrument backlash, so SR_SS_ is difficult to achieve. Comparatively, stress relaxation *sensu lato* (SR_SL_) has practical advantages as it is easier to implement and allows increased precision and sensitivity and, in principle, is independent of mechanical backlash. As we discuss later, each of these tests has a distinct heuristic value.

### Plant material

Experiments were conducted using a diverse set of species growing within the ANU Campus in Canberra, Australia. Most of the experiments were performed at the end of winter and during early spring of 2021; thus, we focused this study on evergreen species. A total of twelve angiosperms and one conifer from ten different families were chosen; the chosen species and the experiments for which they were used are listed in Table S1. As our goal was to test the applicability of uniaxial compression for the measurement of plant water status, replication within a single species was kept at a minimum in an attempt to test the technique in as many species as possible. For experiments in which we used excised tissue, branches were collected using secateurs, placed in a black plastic bag and brought to the lab within five minutes. In the lab, the branches were recut under water and allowed to rehydrate for c. 15 minutes. All experiments were conducted under laboratory conditions (T ∼ 22 °C, RH ∼ 50%).

### Exemplary creep and SR experiments

To illustrate the ability of the instrument to perform the intended techniques and qualitatively assess the equilibration kinetics of leaf squeeze-flow rheometry, we performed creep and SR tests on the lamina (avoiding the midvein and, when possible, secondary veins) of leaves attached to artificially rehydrated branches in two species, *Salvia officinalis* and *Populus nigra*. Leaves from branches collected and treated as described above were pre-compressed under a constant stress of 100 kPa for 5 min to ensure good contact between the leaf surfaces and the compression plates^2^, after which they were subject to: (i) a creep routine at 500 kPa for 5 min followed by recovery at 100 kPa; (ii) a SR_SS_ routine after a 25 µm step followed by recovery after a -25 µm step; and (iii) a SR_SL_ routine after a 25 µm step followed by recovery after a -25 µm step. In all tests, the compression rate during each step was 195 µm s^-1^ and the control loop was operated at 5 Hz.

### Dead leaf tests

The hypothesis that the imposed stress is related to turgor pressure assumes that a significant component of the imposed load is borne by an increased turgor pressure in cells within the compressed region. To verify the role of living tissue in supporting the load as increased turgor pressure, we subjected leaves (N = 3) from a sclerophyllous species (*Quillaja saponaria*) to a SR_SS_ procedure before and after killing the leaf. The choice to perform the test in a sclerophyllous species was made to assess the case where significant resistance to deformation may be offered by the leaf structure even in the absence of turgor. Leaves were pre-compressed under creep at 100 kPa and then compressed with a 25 µm step at a rate of 195 µm s^-1^, followed by relaxation and recovery in periods of 2 min. After recovery, leaves were submerged under water at 80 °C for 30 s; the temperature used for this treatment was chosen according to Yang et al. (2017). Immediately after the 30 s elapsed, leaves were blotted dry and the same routine was repeated. Data from the SR_SS_ experiment were analysed according to Peleg (1979) to quantify the ‘liquidity’ of the sample and the stress relaxation time constant before and after heating. We use the term ‘liquidity’ to quantify the relative stress attained at equilibrium, where a minimum value of 0 means no relaxation (*i*.*e*., an ideal elastic solid) and a maximum value of 1 means full relaxation (*i*.*e*., an ideal liquid) (Peleg, 1979).

### Simultaneous creep and SR sensu lato experiments

Our main hypotheses were that creep can be used to monitor leaf water content, and that SR can be used to quantify changes in leaf turgor pressure. Preliminary experiments indicated that SR_SL_ was a strong predictor of changes in leaf water potential. Thus, capitalising on the simplicity of this method, we first approached the hypotheses by performing simultaneous creep and SR_SL_ experiments.

To perform creep tests, a leaf from an artificially rehydrated branch was compressed in an instrument similar to the one shown in Fig. 2, with the difference that the motor had a planetary gearbox (1:139) and a different load cell (FS2050-1500, TE Connectivity, Switzerland). The gearbox provided increased torque and precision in unidirectional movement, although it increased backlash, reducing the effective resolution of the system due to ‘dead steps’ taken when the motor changed direction. Backlash in this system was *c*. 4 µm. Creep tests were conducted at a pressure of 400 kPa using a microcontroller (Arduino UNO R3, Keyestudio, China) operated with the control loop at 2 H. For the SR_SL_ test, a different leaf from the same branch was compressed using a digital micrometer (SHAHE, China) and the pressure was measured using an amplified load cell (FX29, TE Connectivity, Switzerland) and a 16-bit analog-to-digital converter (ADS1115, Tenstar Robot, China) connected to the microcontroller. Data were logged into a Secure Digital (SD) card. Samples used for SR_SL_ experiments were compressed manually to a pressure of *c*. 800 kPa and left to equilibrate for *c*. 10 min, after which the branch was cut in air to start dehydrating. Branches were dehydrated until the pressure during SR_SL_ reached *c*. 200 kPa, when the branches were recut under water and allowed to rehydrate until they approached equilibrium.

During the experiment we monitored the leaf balancing pressure and the leaf relative water content in three leaves per time point per branch. Excised leaves were placed inside a sealable plastic bag we had previously breathed in (to minimise the vapour pressure deficit therein) and measured within 5 min. The balancing pressure (BP) was measured using a pressure chamber (1505D, PMS Instruments, USA) (Scholander et al., 1965). The leaf relative water content was measured by weighing three leaves and by rehydrating them for 24 h inside a 50 mL plastic tube with the tip of the petiole submerged under water. After, rehydration, leaves were blotted dry with a paper towel, weighed again, and oven-dried at 80 °C to obtain the dry weight. Relative water content was calculated according to Weatherley (1950), as

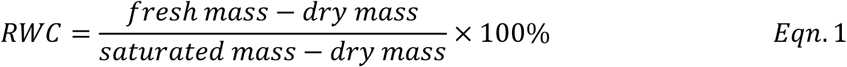

To test the hypothesis that RWC can be approximated as a function of the leaf thickness, we normalised the dimensional compressed leaf thickness during creep h_c_^CR^ relative to its estimated maximum. The maximum h_c_^CR^ was determined as the y-intercept of the regression between h_c_^CR^ and BP considering data where this relation was linear (*i*.*e*., near ‘full hydration’). The relative compressed leaf thickness during creep (H_c_^CR^) was obtained as

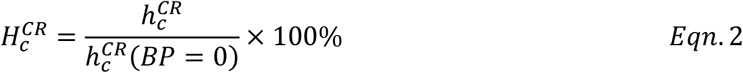

### Pressure-volume curves

Leaves from each branch used for the creep and SR_SL_ experiment were used to build pressure-volume curves. Before cutting the branch in air to start dehydration, three leaves were excised and placed inside a sealable plastic bag we had previously breathed in. Leaves were then subject to alternate measurements of balancing pressure and fresh mass until the leaves looked wilted. Leaves were oven-dried for 48 h at 80 °C to obtain the dry weight.

Leaf pressure-volume curves were analysed by plotting the inverse of the leaf balancing pressure (1/BP) against the leaf fresh mass (Tyree and Hammel, 1972). The linear portion of the data was fitted with a linear regression and the TLP was determined as the highest measured fresh mass within the fitted data. The leaf water volume was normalised relative to the volume at the turgor-loss point as

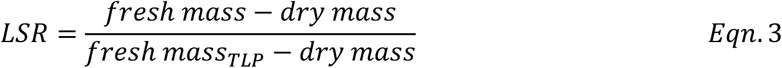

where LSR stands for the ‘leaf swelling ratio’. Following determination of LSR, 1/BP was plotted against LSR and BP_TLP_ was obtained as the inverse of the regression function evaluated at LSR = 1. The leaf apoplastic water fraction (af) was obtained as the x-intercept between LSR and 1/BP. Using the linear regressions obtained via this method, an estimate of the osmotic pressure Π at all levels of hydration was obtained, whereby the turgor pressure was derived as P = Π – BP. The leaf bulk elastic modulus (E) was obtained by linear regression, in the positive turgor range, as E = ΔP/ΔLSR.

Data from leaf pressure-volume curves were used to test the hypothesis that P can be estimated from stress relaxation. To do this, a linear regression equation was used to obtain P as a function of BP; these regressions (all with R^2^ > 0.99) were then used to estimate the turgor pressure from the BP measured in the samples used for SR_SL_.

### In vivo measurements

*Avicennia marina* subsp. *australasica* plants were grown from propagules collected in November 2019 along the Clyde River estuary, NSW, Australia. Propagules were germinated in sand saturated with a 50% seawater solution. After depletion of cotyledonary reserves, plants were transplanted into 10 L plastic buckets with a 50% seawater solution and fertilised occasionally with a liquid seaweed fertiliser (Seasol, Seasol International, Australia). At the time of the experiment, each bucket contained six plants bearing c. 10 mature leaf pairs. Water was not added during the experiment. A full-spectrum LED light source (TS1000, Mars Hydro, China) installed c. 50 cm above the canopy provided a 14/10 h light/darkness cycle. Creep and SR_SL_ tests were conducted as described for the simultaneous creep and SR_SL_ experiments using the instrument shown in Fig. 2, with the difference that creep tests were conducted at 100 kPa.

### Data analyses

Data were curated in MS Excel and analysed in R version 4.0.5. All data presented are calculated from raw measured values, except for those measured in *Avicennia marina*, which were filtered using a third-degree polynomial moving average with a 1-min window (Savitzky and Golay, 1964) using the ‘sgolayfilt’ function of the ‘signal’ package. Regressions were performed using the built-in ‘lm’ function and the ‘nlme’ package and assessed via visual inspection of residual distributions. Figures were built using the ‘ggplot’ package and stylised in Adobe Illustrator (Adobe Systems, USA).

## Results

### Instrument performance

The custom-built leaf squeeze-flow rheometer performed well in calibration tests. We found good agreement between the stepper motor instruction and the micrometer reading (Fig. 2C). The observed error (RMSE = 1.1 µm) was associated with the system backlash, which was *c*. 2 µm (Fig. S1). When conducting creep tests on an empty sample (Fig. 2A, D), the system operated as instructed (Fig. 2B), although stress overshoot and undershoot were observed immediately after changing the input stress in each step (Fig. 2A). The system deflection under load was predicted well by a third-order polynomial function, displaying no hysteresis (Fig. 2E), which allowed accurate estimation of the system dimensions for any given stress. The instrument was found to be temperature-sensitive (Fig. 2F). A thermal cycle between 5 °C and 40 °C indicated that the system deflection increased approximately logarithmically with temperature, being most sensitive (*c*. 30 nm K^-1^) at low temperatures.

### Exemplary curves and dead leaf compression test

Leaves from two species (*Populus nigra* and *Salvia officinalis*) pre-compressed under a stress of 100 kPa provided example time-dependent responses to leaf uniaxial compression. In all tests, we observed characteristic times of a few minutes (Fig. 3). In the case of creep tests (Fig. 3A, B), *Salvia* leaves compressed to a lower relative thickness than *Populus* leaves; similarly, stress relaxation *sensu stricto* (Fig. 3C, D) and *sensu lato* (Fig. 3E, F) showed that the equilibrium stress reached after a 25 µm compression step was higher in the case of *Populus*. The technique consistently revealed mechanical differences between the species. While SR_SL_ and SR_SS_ yielded similar results, SR_SS_ produced noisier data in the case of poplar (Fig. 3C, E). In the SR_SS_ procedure, the sample thickness was maintained constant to about ± 0.5 µm.

**Figure 3.**
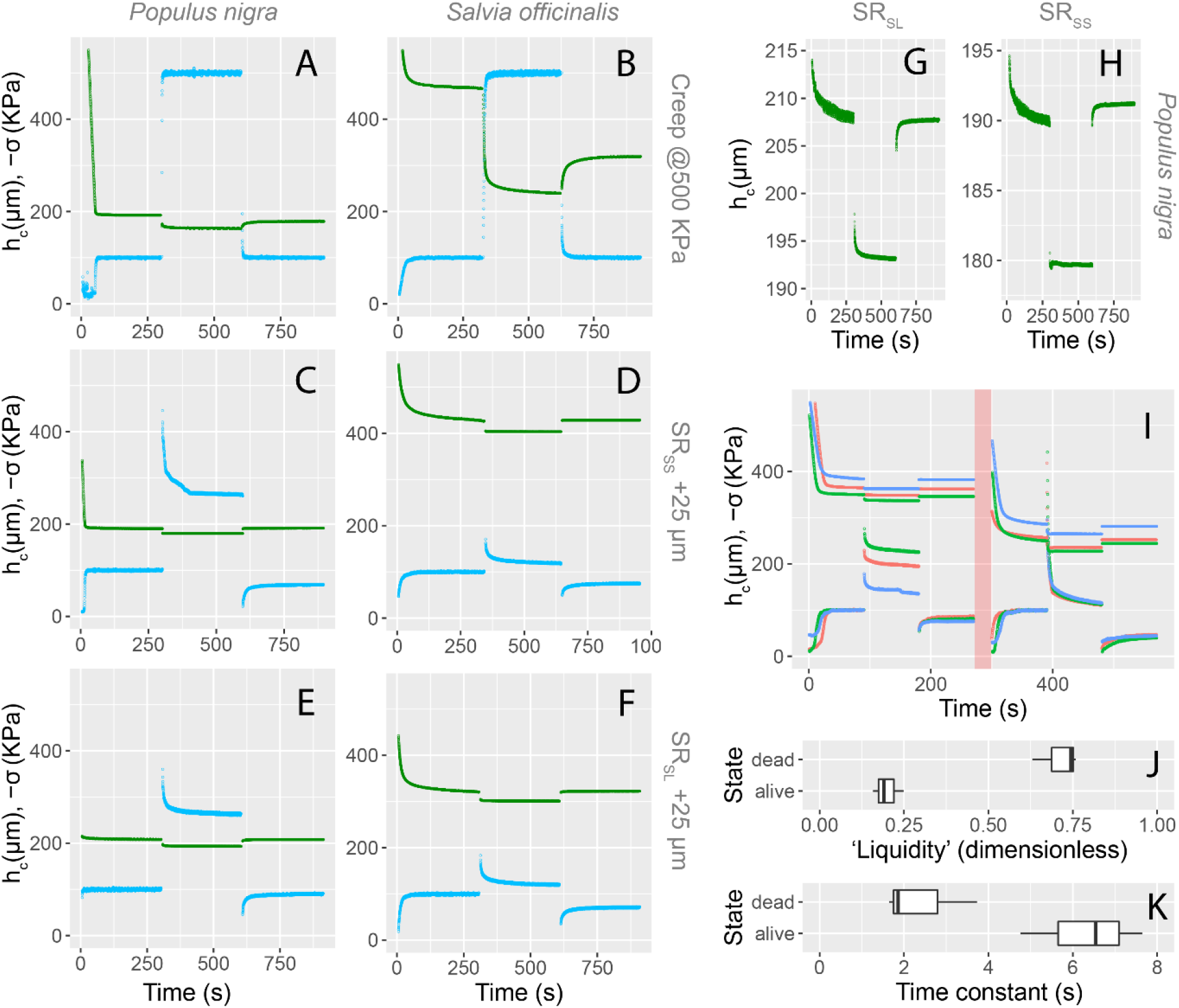
A-F: Examples of creep and stress relaxation tests. Leaves were pre-compressed under a stress of 100 kPa to ensure good contact between the leaf surfaces and the compression plates. Leaves were then subject to: creep at 500 kPa (A, B); stress relaxation *sensu stricto* (SR_SS_) (i.e., with the spindle position adjusted to maintain constant sample thickness) following a ± 25 µm step at 195 µm s^-1^ (C, D); and stress relaxation *sensu lato* (SR_SL_) (i.e., with the spindle position fixed) following a ± 25 µm step at 195 µm s^-1^ (E, F). Green curves correspond to the sample thickness; blue curves correspond to the applied compressive stress. These tests were performed on living leaves attached to a branch with the cut end maintained under water, so the water status of the (uncompressed) tissue is assumed constant. N = 1. G, H: sample thickness obtained during the stress relaxation *sensu lato* and *sensu stricto* tests shown in panels C and E. I: Stress relaxation *sensu stricto* tests on three *Quillaja saponaria* leaves before and after immersion under water at 80 °C for 30 s (red shaded area); colours denote different samples. Samples were subjected to creep at 100 kPa followed by stress relaxation and recovery after a ± 25 µm step at 195 µm s^-1^. Upper curves correspond to the sample thickness and lower curves correspond to the applied compressive stress. Note the difference in the leaf thickness and relaxation kinetics before and after killing the tissue. J: Fitted coefficients using the method by Peleg (1979) to quantify the ‘liquidity’ of the samples, where a value of zero means no relaxation (*i*.*e*., an elastic solid) and a value of one means complete relaxation (*i*.*e*., a liquid); *P* = 0.0005 (Student’s t-test). K: Stress relaxation half-time according to the method by Peleg (1979); *P* = 0.022 (Student’s t-test). N = 3.

Compressing leaves from the sclerophyllous species *Quillaja saponaria* revealed changes in the leaf stress relaxation kinetics following heat damage (Fig. 3I). After submerging tissues under 80 °C water for 30 s, the ‘liquidity’ of the samples increased from 0.199 ± 0.026 to 0.711 ± 0.041 (mean ± SE) (*t* = -10.54; *P* = 0.0005) (Fig. 3J); the time half-time of stress relaxation decreased from 6.32 ± 0.84 s to 2.41 ± 0.66 s (*t* = 3.65; *P* = 0.022) (Fig. 3K).

### Creep and stress relaxation as guiding principles to monitor leaf water status

The leaf thickness under creep (h_c_^CR^) and the compressive stress applied during stress relaxation *sensu lato* (−σ^SRSL^) were measured during a dehydration-rehydration cycle. Cutting in air induced a decline in h_c_^CR^ and −σ^SRSL^, which was largely reverted by cutting the stems under water. Stress relaxation curves during dehydration were varied in shape and can be described as S-shaped or J-shaped. For our experimental conditions, the time required for a 50% decrease in −σ^SRSL^ since the time of cutting in air varied widely between species, with a mean ± SE of 2.1 ± 0.7 h. In all species, the recovered −σ^SRSL^ was lower than the initial −σ^SRSL^; a similar pattern was observed for h_c_^CR^, with the exception of *Corymbia citriodora* (Fig. 4 B6). Variations to typical rehydration kinetics curves were sometimes observed, *e*.*g*., in a *Callistemon viminalis* branch which exhibited an unusual rehydration pattern (Fig. 5A). The relation between h_c_^CR^ and −σ^SRSL^ was shown to be non-linear; in most cases, the slope of –h_c_^CR^ /σ^SRSL^ increased during dehydration and also during rehydration. The latter steepening of the curves resulted in hysteresis loops which displayed wide variation in terms of area and recovery.

**Figure 4.**
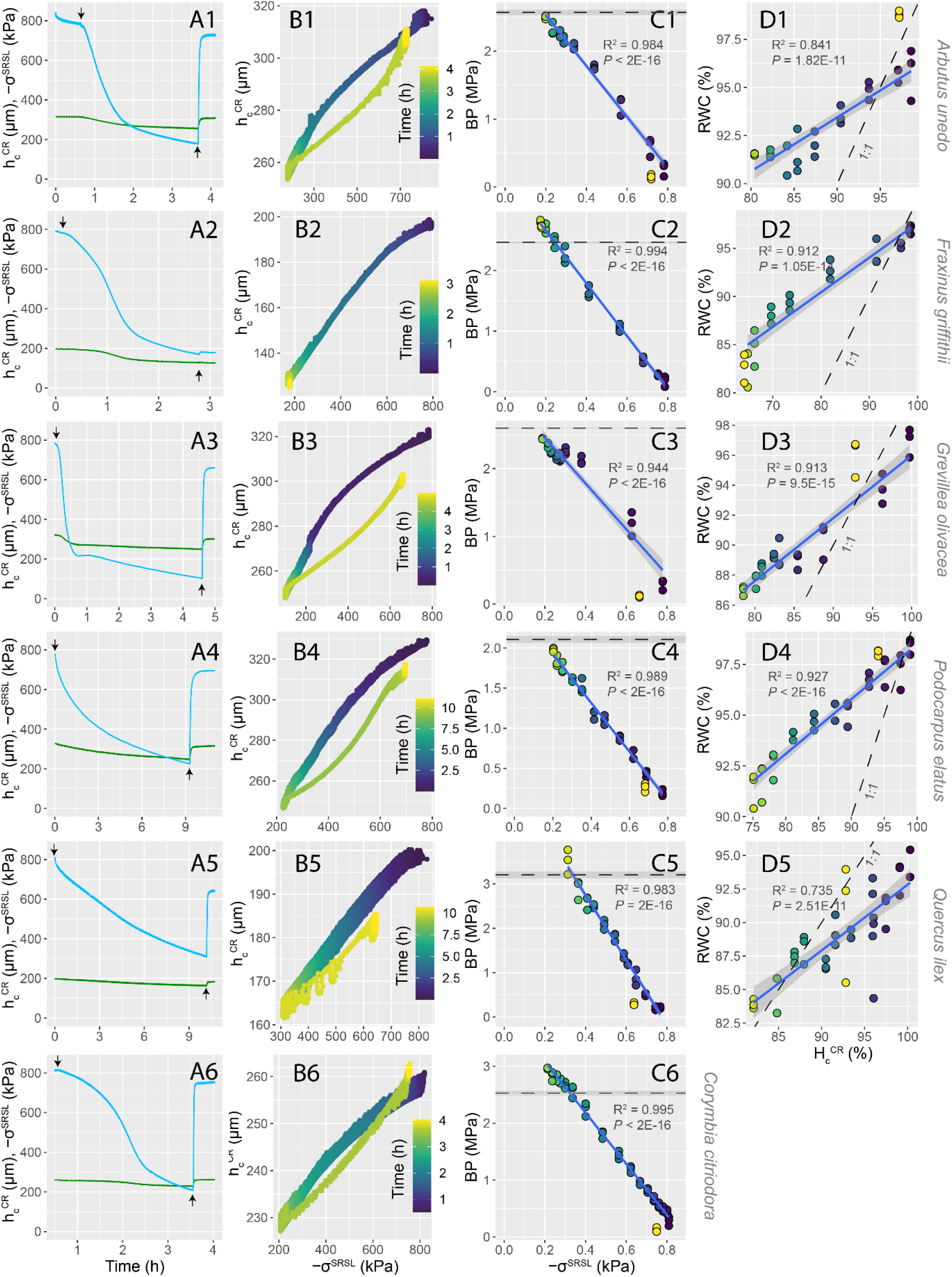
Simultaneous creep and stress relaxation *sensu lato* (SR_SL_) experiments performed on a single branch per species under changing water status. A1-A6: Leaf thickness measured during creep at 400 kPa (h_c_^CR^, green lines) and stress applied during SR_SL_ (−σ^SRSL^, blue lines) for different species during a dehydration-rehydration cycle. Down and up arrows indicate the time at which branches were cut in air and under water to start dehydration and rehydration, respectively. B1-B6: Relation between the thickness measured during creep and the stress measured during SR_SL_. C1-C6: Relation between the balancing pressure measured in three leaves per branch per time point and the stress measured during SR_SL_. The horizontal dashed lines indicate the balancing pressure at the turgor-loss point (BP_TLP_); shaded areas correspond to standard error. D1-D5: Relation between the leaf relative water content (RWC) measured in three leaves per time point and the relative leaf thickness measured during creep (H_c_^CR^); dashed lines correspond to the 1:1 line. Regressions in columns C and D only consider points obtained during dehydration. Colour scales for time shown in column B also apply to panels in columns C and D within the same row. N = 1.

**Figure 5.**
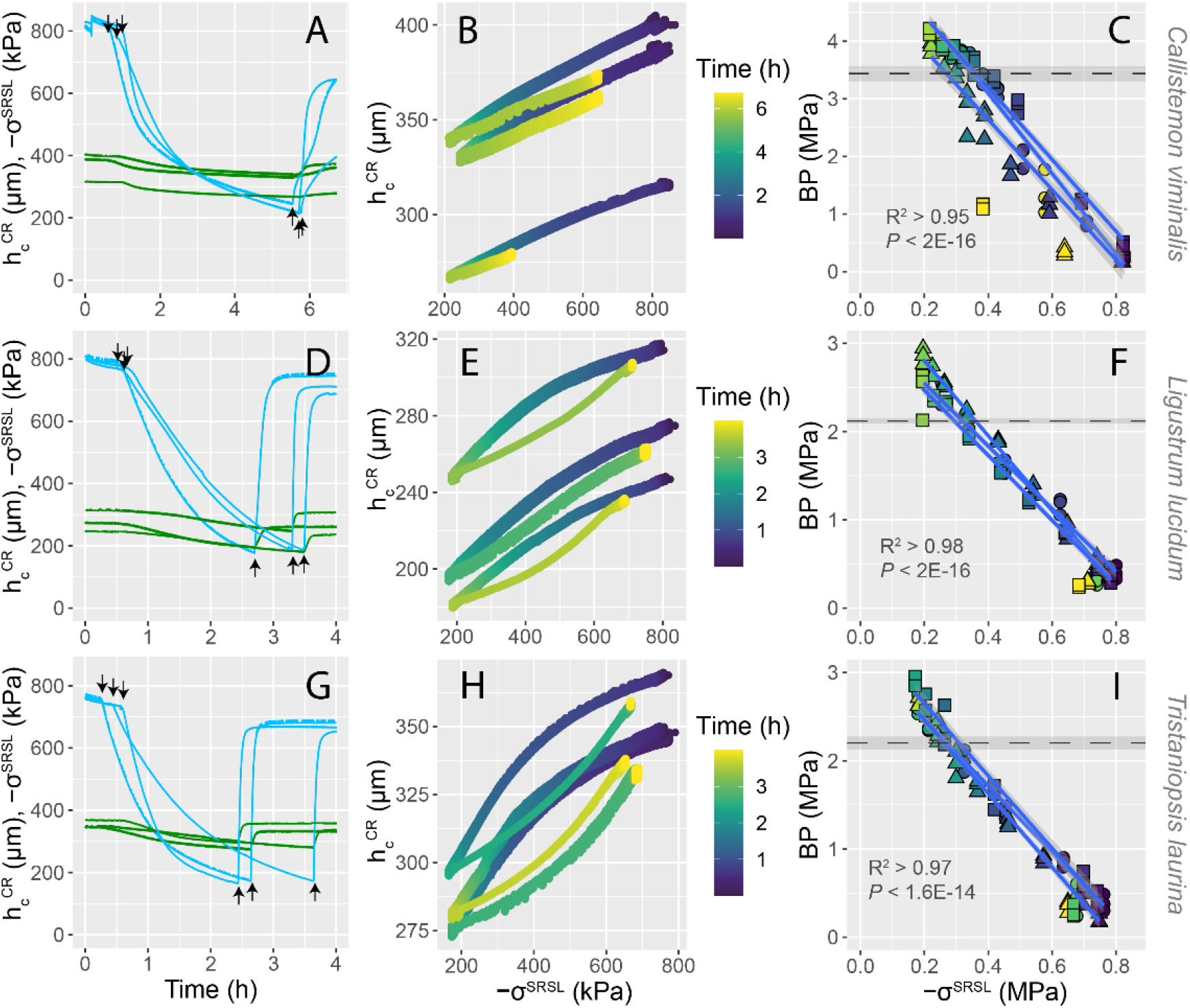
Simultaneous creep and stress relaxation *sensu lato* (SR_SL_) experiments performed on three branches per species under changing water status. A, D, G: Leaf thickness measured during creep at 400 kPa (h_c_^CR^, green lines) and stress applied during SR_SL_ (−σ^SRSL^, blue lines) for different species during a dehydration-rehydration cycle. Down and up arrows indicate the time at which branches were cut in air and under water to start dehydration and rehydration, respectively. B, E, H: Relation between the thickness measured during creep and the stress measured during SR_SL_. C, F, I: Relation between the balancing pressure measured in three leaves per branch per time point and the stress measured during SR_SL_. The horizontal dashed lines indicate the balancing pressure at the turgor-loss point (BP_TLP_); shaded areas correspond to standard error. Regressions only consider points obtained during dehydration. Colour scales for time shown in B, E and H also apply to panels C, F and I, respectively. N = 3.

Stress relaxation *sensu lato* provided a good estimator of leaf balancing pressure (BP) and the relation was, in most cases, approximately linear (Figs. 4 & 5). The mean ± SD R^2^ value for the linear regressions between −σ^SRSL^ and BP was 0.98 ± 0.01 (N = 15 branches from 9 species). Thus, the system demonstrated good capability to estimate BP non-invasively. The leaf relative thickness measured during creep (H_c_^CR^) was positively related to the leaf relative water content (RWC). The mean ± SD R^2^ value for the linear regressions between H_c_^CR^ and RWC was 0.87 ± 0.08 (N = 5 branches from 5 species). In some species, however, the relation between H_c_^CR^ and RWC was clearly non-linear (e.g., in *Fraxinus griffithii*, Fig. 4 D2). Overall, the technique was less accurate in estimating RWC than BP.

### Relation to leaf pressure-volume parameters

The parameters extracted from the relation between the applied stress (−σ^SRSL^) and BP during stress relaxation (Figs. 4 & 5) and those obtained between leaf relative thickness (H_c_^CR^) and RWC (Fig. 4) were related to leaf pressure-volume parameters. In the case of the regression parameters between −σ^SRSL^ and BP, the intercept at −σ^SRSL^ = 0 and the slope were correlated with the osmotic pressure at the turgor-loss point (Fig. 6A, B) and the leaf bulk elastic modulus (Fig. 6C, D); no significant relation was found between these parameters and the leaf apoplastic fraction (Fig. 6E, F). In the case of the regression parameters between H_c_^CR^ and RWC, the intercept at H_c_^CR^ = 0 and the slope were related to the osmotic pressure at the turgor-loss point (Fig. 6G, H); no significant relation between intercept and slope between H_c_^CR^ and RWC and the leaf bulk elastic modulus or the leaf apoplastic water fraction was found (Fig. 6 I-L). We note that the results for the relationships between H ^CR^ and RWC may be weaker due to a smaller sample size.

**Figure 6.**
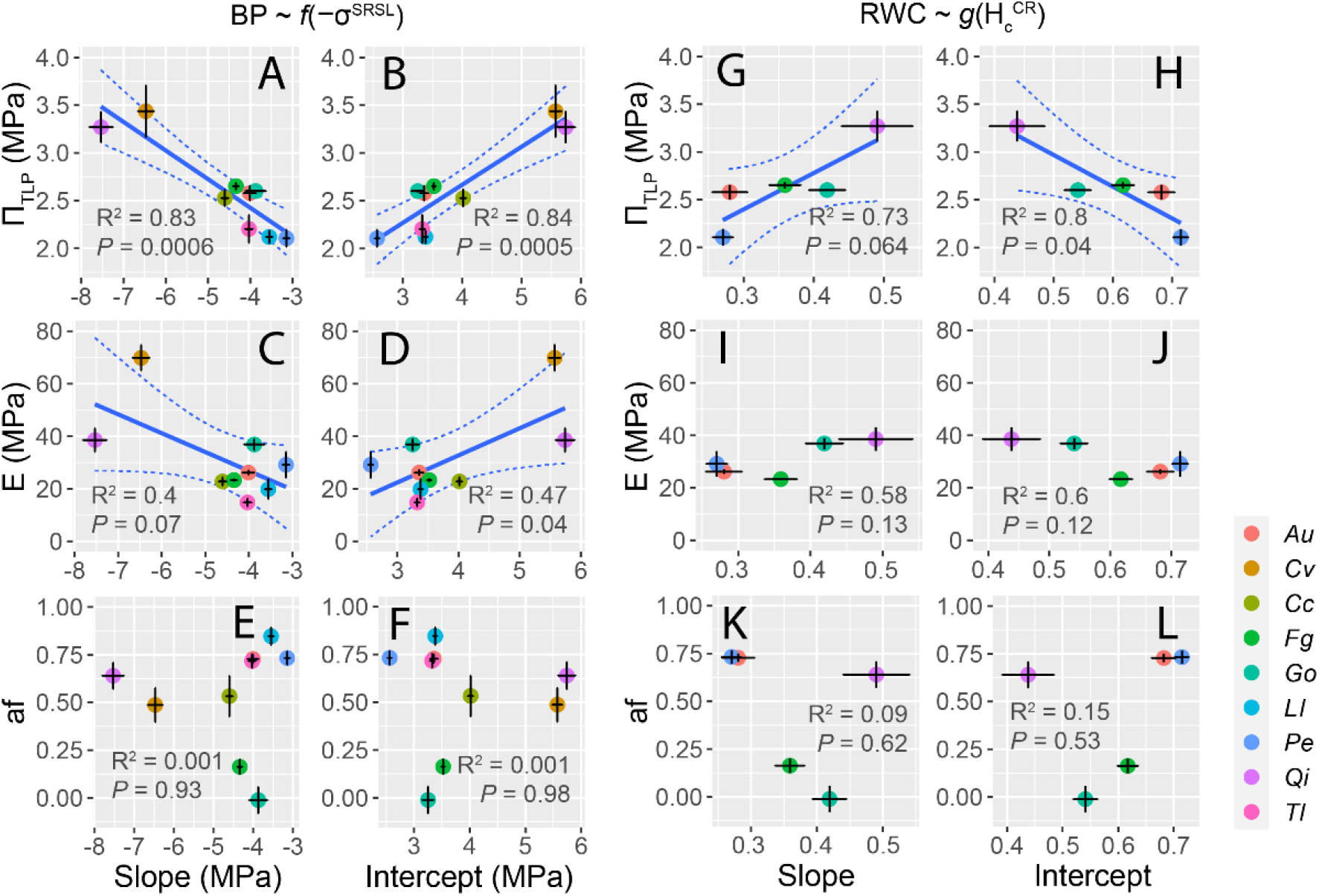
Relation between pressure-volume parameters and the linear regression coefficients from the experiments shown in Figs. 4 and 5. A-F: Relation between the regression coefficients of the stress applied during stress relaxation *sensu lato* (−σ^SRSL^) and leaf pressure-volume parameters. G-L: Relation between the regression coefficients of the leaf relative thickness measured during creep (H_c_^CR^) and leaf pressure-volume parameters. Colours denote different species as per the legend. *Au* = *Arbutus unedo*; *Cv* = *Callistemon viminalis*; *Cc* = *Corymbia citriodora*; *Fg* = *Fraxinus griffithii*; *Go* = *Grevillea olivacea*; *Ll* = *Ligustrum lucidum*; *Pe* = *Podocarpus elatus*; *Qi* = *Quercus ilex*; *Tl* = *Tristaniopsis laurina*. Π_TLP_ = leaf osmotic pressure at the turgor-loss point; E = leaf bulk elastic modulus; af = leaf apoplastic water fraction.

Balancing pressure measurements shown in Figs. 4 and 5 were used to estimate the leaf turgor pressure (P) during dehydration using data from leaf pressure-volume curves. As expected from the shown relation between BP and –σ^SRSL^, linear regression showed that –σ^SRSL^ was strongly correlated with leaf turgor pressure (Fig. 7), with an R^2^ > 0.95 across 15 branches from 9 different species.

**Figure 7.**
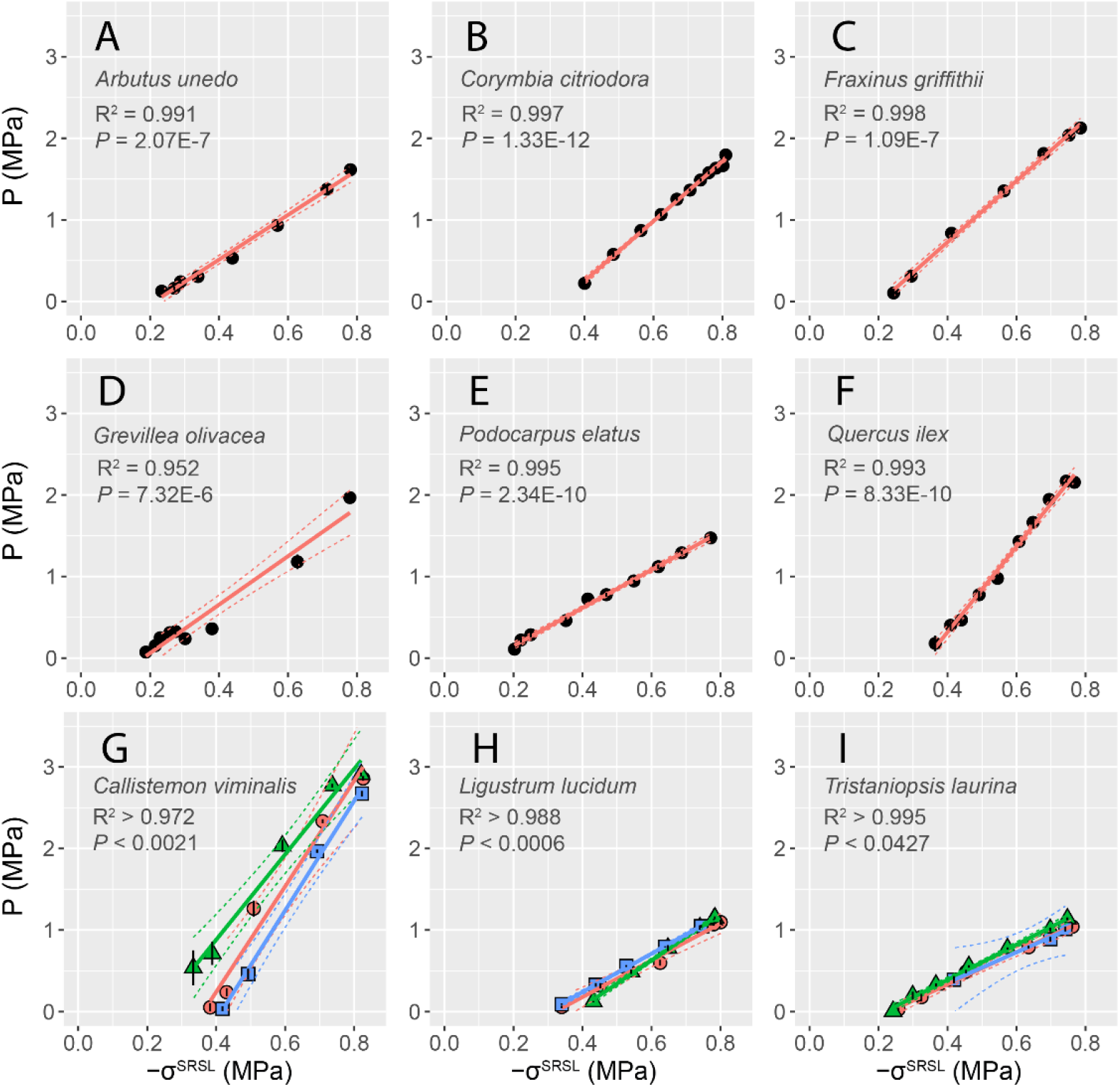
Relation between the stress applied during stress relaxation *sensu lato* (−σ^SRSL^) and the turgor pressure (P) estimated from leaf pressure-volume curves. Colours and symbols in panels G-I denote different plants.

### In vivo measurements

Measurements performed on a living potted mangrove (*Avicennia marina* subsp. *australasica*) revealed that the technique was capable of measuring small changes in water status induced by light (Fig. 8). As expected, thickness and pressure decreased during the day and increased during the night. Immediately after light exposure, the plant exhibited sudden increases in pressure followed by a sharp decline. Over the four days, we observed a *c*. 7 µm decrease in leaf thickness and a *c*. 20 kPa decrease in the applied stress. Diurnal variations in h_c_^CR^ were in the order of < 5 µm; daily variations in −σ^SRSL^ were < 35 kPa. These small changes were not well resolved by raw measurements, but implementation of a Savitzky-Golay filter with a 1-min window produced data with acceptable resolution. Dimensional changes due to variations in the micrometer temperature were negligible in this experiment (∼0.075 µm expansion/contraction during light and dark periods, according to the relation shown in Fig. 2G).

**Figure 8.**
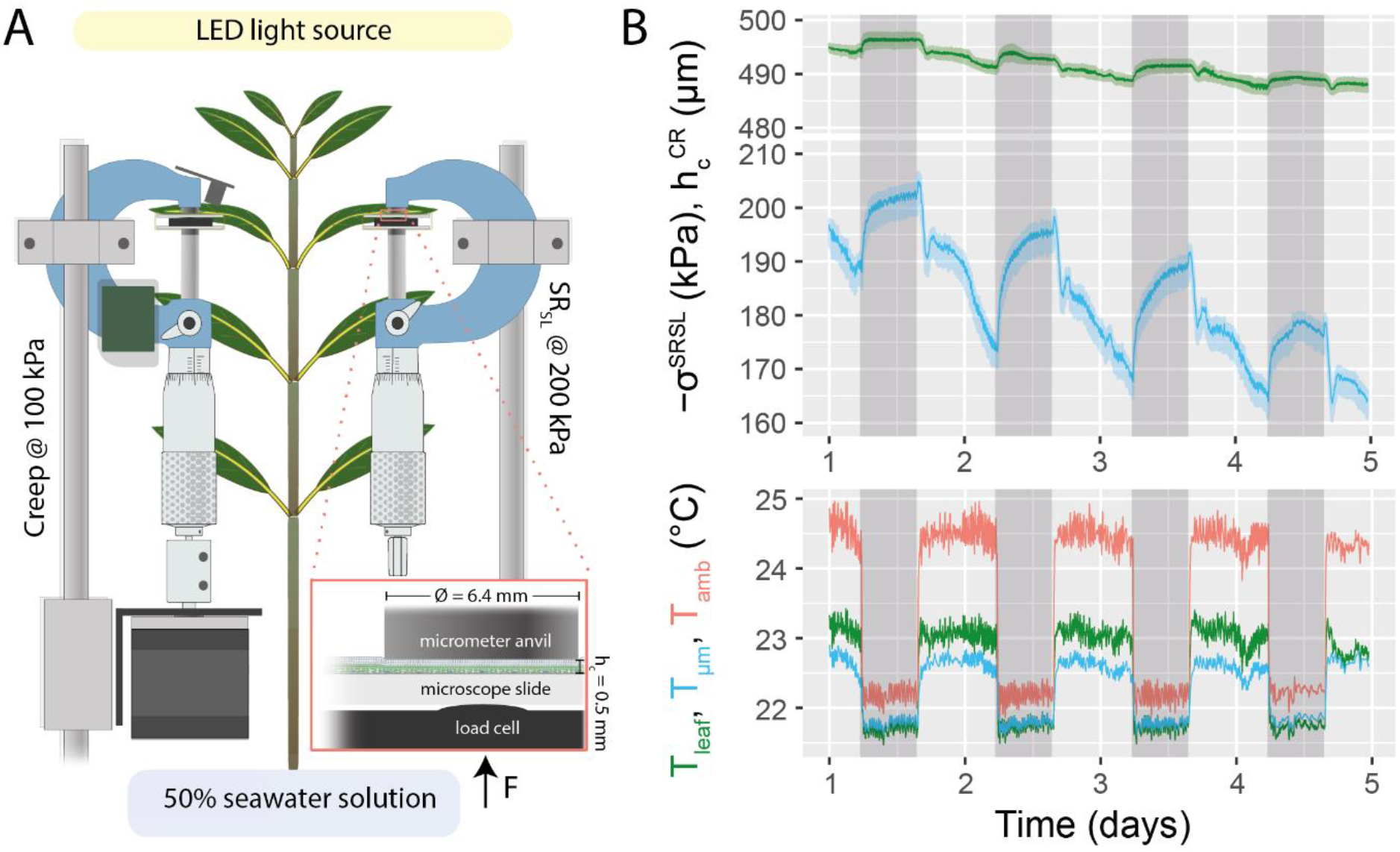
Simultaneous creep and stress relaxation *sensu lato* experiments performed on a living potted plant (*Avicennia marina* subsp. *australasica*) under laboratory conditions. A: Simplified view of the experimental setup (electrical connections are omitted for simplicity). A motorised micrometer was used for running a creep test at 100 kPa and a manual micrometer was used to perform a stress relaxation *sensu lato* test at *c*. 200 kPa. The plant was maintained inside a tub with a 50% seawater solution and was irradiated using a full-spectrum LED light source. The red rectangular area shows a magnified view of an idealised uniaxially compressed *Avicennia* leaf (to scale). B: Results from the simultaneous creep and SR_SL_ experiment. Shaded grey bars indicate periods of darkness. These measurements were filtered using a third-degree polynomial moving average (Savitzky and Golay, 1964) with a 1 min window; shaded coloured bands correspond to the raw measurements. Temperature-induced variations in the system thickness calculated from Fig. 2G are negligible in this experiment (*c*. 75 nm variation between during day and night). The small and steady decline in thickness and mechanical stress may be due to an increase in salinity of the hydroponic solution. T_leaf_ = leaf temperature; T_µm_ = micrometer frame temperature; T_amb_ = temperature of the infrared sensor directly above the leaf. N = 1.

## Discussion

Leaf squeeze-flow rheometry was applied as a method to monitor plant water status. The stress measured during stress relaxation *sensu lato* was related to turgor pressure (R^2^ > 0.95) and leaf water potential (measured as −BP) (R^2^ > 0.94); the relative leaf thickness measured during creep was related, although sometimes non-linearly, to leaf relative water content (R^2^ > 0.74). In its current implementation, squeeze-flow rheometry enabled reasonably precise non-destructive measurement of plant water status with high temporal resolution at low cost, opening new opportunities for the measurement of plant hydration dynamics.

### Instrument performance

The leaf squeeze-flow rheometer performed adequately; given its cost and simplicity, it may be useful to plant scientists interested in studying water in plants. As the application of this technique is novel to the study of plant hydration dynamics, we briefly discuss key aspects for future improvement.

Mechanical backlash was the main limitation to the positioning accuracy of the system, and the main cause of the inability to perform SR_SS_ experiments when the direction of stress relaxation changes. The mechanical backlash may originate from gaps in the micrometer screw or the linear guide. An effective solution to this problem could be the incorporation of a linear-variable differential transformer (LVDT), which can provide absolute positioning at sub-micron resolution. Despite this current limitation, the observed precision of the instrument compared favourably to commercially available systems to measure leaf thickness. Therefore, we found the instrument acceptable for the intended purpose.

Exemplary creep and stress relaxation tests (Fig. 3) showed that the instrument was capable of performing the tests intended, with two main limitations. First, SR_SS_ tests were performed adequately only in the cases where the direction of stress relaxation does not change. Second, the time required to achieve a constant stress during creep was dependent on the stiffness of the sample, with softer samples (e.g., *Salvia officinalis*) taking longer to reach a constant stress. Currently, the instrument only incorporates proportional control of the spindle position; use of more advanced control systems, such as proportional-integral-derivative (PID) control, can be implemented to achieve better performance during creep tests.

The instrument exhibited a temperature sensitivity of order 30 nm K^-1^ and is presently only suited for measurements at constant or nearly constant temperature. In the case of creep experiments, the estimated load cell deflection can be simply subtracted from the actual measurements; however, during stress relaxation, the micrometer position must be adjusted to maintain a constant sample thickness. Temperature-corrected positioning control and/or use of dimensionally stable composites may be employed to correct this issue.

### Time-dependent behaviour during leaf squeeze-flow rheometry

In this study we emphasized the study of leaf squeeze-flow rheometry under changing water status when the compressed region was in hydrostatic equilibrium. However, results from few example curves comparing two species with distinct leaves, poplar (*Populus nigra*) and sage (*Salvia officinalis*), revealed rheological differences between the species during creep and SR. In poplar, we observed nearly complete recovery of the original thickness or pressure after undergoing creep and stress relaxation tests; by contrast, sage leaves displayed large irreversible losses of thickness, particularly during the creep test. Although we did not investigate the physical basis of creep and SR, it is likely that these differences are determined by the diffusion of water out of the compressed cells, loss of intracellular solute due to compression, and/or plastic yield of the tissue. These phenomena are interesting in their own right and may be explored further. As this is, to our knowledge, the first approximation to leaf squeeze-flow rheometry, we briefly describe the effects of leaf uniaxial compression on the components of plant water status.

Leaves with flat laminae can be described as a cellular composite in the form of a sandwich beam (Gibson et al., 1988). Tissue on the sunny side of the leaf is packed with cylindrical cells with their long axis perpendicular to the epidermis, forming a layer known as the ‘palisade’ parenchyma; the shady side of the leaf can be described as an open-cell foam and is aptly called ‘spongy’ parenchyma (Borsuk et al., 2022). Most leaf tissues are porous, so uniaxial compression can presumably deform cells by changing their shape, *i*.*e*., by making them wider and shorter. An idealised average-cell view of the time-dependent changes in water status that occur with uniaxial compression is shown in Fig. 9 (here we illustrate the case of stress relaxation, but a similar behaviour would be expected during creep). This simplified view excludes non-ideal osmotic behaviour and plastic yield, so may only be applied to cells experiencing small strains. In a timescale of minutes, relaxation of tension in the cell wall and water diffusion out of the cell cause the cell hydrostatic pressure to drop to *P* + *ΔP*_*c*_. Equilibrium (3) is reached when the new hydrostatic pressure is balanced by the increased osmotic pressure *Π*_*c*_.

**Figure 9.**
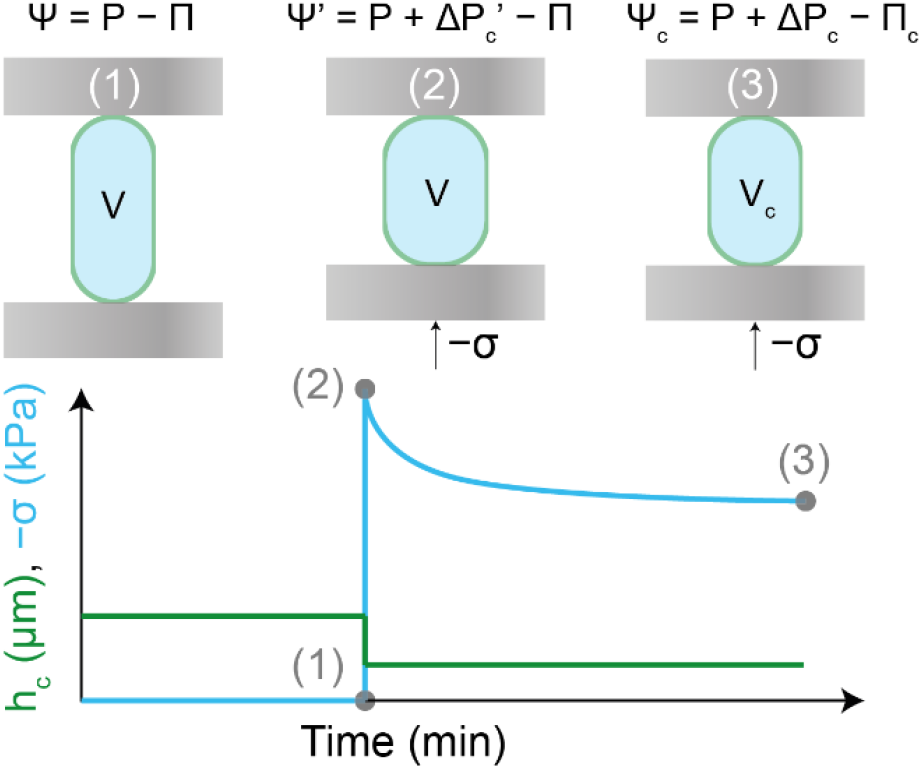
Idealised average-cell view of the changes in leaf water status brought by leaf uniaxial compression during stress relaxation. An uncompressed cell (1) is quickly compressed between two parallel plates, inducing immediate cell deformation with no change in the cell volume V, transiently raising the cell hydrostatic pressure to *P* + *ΔP*_*c*_*′* (2) ^3^. The plates are then maintained at a fixed position.

Under the mechanical stresses we worked with, water is effectively incompressible. Therefore, immediately after loading, we expect the volume of the cells to remain unchanged, and the change in shape to induce an increased tension in the cell walls, increasing the intracellular hydrostatic pressure to (*P* + *ΔP*_*c*_*′*), where the last term indicates the maximum change in intracellular hydrostatic pressure caused by compression. In this state, the water potential of the compressed cells is 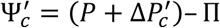, while outside the compressed region it is simply *Ψ* = *P*− *Π*. Water within the compressed region may equilibrate with water outside the compressed region, and two processes may be involved in this relaxation: first, water from inside the cell has to diffuse across the cell membrane into the cell wall; and second, water has to diffuse via the apoplastic space to the outside tissue. If membrane diffusion represents the only resistance to water movement, then the water potential in the compressed apoplast may be assumed to remain equal to Ψ (*i*.*e*., the apoplast remains in equilibrium with the tissue outside); if apoplastic diffusion represents the only resistance to water movement, then the water potential in the compressed apoplast is likely to transiently approach zero as free water accumulates in the apoplastic space, before dissipating along a pressure gradient. Probably, both of these resistances to water movement play a role in determining the kinetics of stress relaxation, so the water potential outside the cells during the equilibration process is unknown. Therefore, here we restrict the discussion to the initial and final states.

If the volume of the compressed tissue is negligible in relation to the volume of the tissue studied (*e*.*g*., the whole leaf, branch or plant), the water potential of the uncompressed tissue may be assumed constant. Thus, if the compressed region equilibrates with water outside the compressed region, then water redistribution is expected to lower *ΔP*_*c*_^*′*^ and raise the osmotic pressure in cells within the compressed region, until they reach equilibrium values *ΔP*_*c*_ and *Π*_*c*_. This can be written as

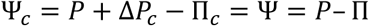

so *ΔP*_*c*_ = *Π*_*c*_− *Π*. Making the approximation that the intracellular osmotically inactive volume is negligible and the cell osmotic behaviour is ideal, then a cell having *n* osmoles of solute and an original volume *V* shrinks to *V*_*c*_, where *ΠV* = *nRT* = *Π*_*c*_*V*_*c*_. Defining the volumetric strain as *v* = (*V*_*c*_ − *V*)/*V* (which is negative for compression) and combining these equations gives

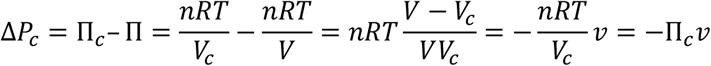

From the Boyle-Van’t Hoff relation and from the definition of the volumetric strain, it follows that *Π*_*c*_ ≅ *Π*/(1 + *v*) (Philip, 1958), so

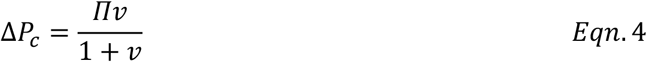

From the point of view of the cell symplast, Eqn. 4 could be viewed as a constitutive relation. However, *ΔP*_*c*_ and *v* are not readily measurable, and the relation between −*σ* and *ΔP*_*c*_ is not simple. While we do not know how the imposed stress relates to the change in turgor, it may be reasonable to make the approximation that they are proportional, given the strong linear relationships found between −σ^SRSL^ and P (Fig. 7). Thus, −*σ ∝ Πv*/(1 + *v*) (for small strains, *Π*_*c*_ ≈*Π*, so the compressed cell may be approximated as linear-elastic). Further studies are needed to compare this simplified average-cell model to more detailed models of uniaxially compressed leaf tissues, which may include anatomy and cellular inhomogeneity.

### Loading of dead leaves

Loading living and dead leaves from the sclerophyllous species *Quillaja saponaria* showed that stress-relaxation parameters were significantly affected by heat-induced cell lysis. The marked increase in the ‘liquidity’ of the samples is consistent with expectation that much of the mechanical load in the living leaves was supported by turgor. Of course, dead leaves did not behave like an ideal liquid either, because some of the load would have been supported by the leaf structure. This effect is likely to become greater as tissue density increases (and porosity decreases) during loading (Gibson et al., 1982). Heat-induced lysis also decreased the stress relaxation half-time. The faster equilibration of dead samples may be due to the removal of the cell membrane, which reduces the resistance to water movement out of the cells; however, it is also likely that the rheology of the leaf is affected due to release of cellular materials that may increase the viscosity of the medium. We note that the change in the SR half-time should be considered qualitatively, as dead leaves shrunk considerably after lysis and thus flow in the dead samples occurred through a smaller leaf cross-sectional area.

### Creep and stress relaxation as measurement paradigms to monitor leaf water status

Creep and stress relaxation are routine tests used in rheology to study the deformation and flow kinetics of viscoelastic specimens. Although they are widely applied in material sciences, these measurement paradigms have not, to our knowledge, been applied to living plant tissues and their potential value in monitoring tissue water status *in vivo* has not been recognised. Given their simplicity and precision, these tests might become useful techniques to monitor plant hydration dynamics.

Experimental evidence indicated that the applied stress measured during stress relaxation *sensu lato* (−σ^SRSL^) can be reliably used as an indicator of changes in leaf turgor pressure. It must be remembered, however, that −σ^SRSL^ is not a direct measurement of P. Possibly due to complex changes in cell wall geometry and stresses, the slopes of P vs −σ^SRSL^ were greater than unity by a factor of several (Fig. 7). The relation between σ^SRSL^ and BP was also approximately linear (Figs. 4 & 5) and their relation varied according to Π_TLP_ (Fig. 6). Although the predictive power of −σ^SRSL^ was high, residual distributions indicate that its relation to BP is not always linear over the range studied. In some species, such as *Callistemon viminalis* and *Corymbia citriodora*, we observed a slight change in slope around the TLP, whereas in *Ligustrum lucidum, Tristianopsis laurina* and *Fraxinus griffithii* the relation was approximately linear beyond the TLP. In other species, we did not reach dehydration levels beyond the TLP. Departures from linearity may be caused by steeper changes in the leaf osmotic pressure once the tissues become flaccid. Other departures from linearity in the relation between BP and −σ^SRSL^ may have been associated to a lack of hydrostatic equilibrium across the plasma membrane. For example, in *Grevillea olivacea*, most turgor was lost within the first 30 min since cutting in air, during which the observed BP was higher than predicted by −σ^SRSL^ (Fig. 4 C3). The latter may suggest the tension of xylem water to be increasing at a higher rate than the reduction in cell turgor. Thus, the simplified proportional relation between −σ^SRSL^ and BP may be restricted to cases where equilibrium across the plasma membrane is met.

As leaf thickness is not constant during SR_SL_, the relation between −σ^SRSL^ and BP is to some extent dependent on the mechanical compliance of the measurement system (Fig. 2E). Thus, our results are restricted to the instrument we used, or one with comparable mechanical compliance, and it remains to show the relation between −σ^SRSS^ and BP. While we are unable to perform SR_SS_ tests under changing water status in our custom-built squeeze-flow rheometer, researchers with access to universal testing machines or dynamic mechanical analysers could test this relationship in a matter of hours. Thus, while SR_SL_ has advantages due to the simplicity of the instrument and reduced noise in the data, SR_SS_ may provide greater replicability as the results should be independent of the instrument’s mechanical compliance.

The relation between H_c_^CR^ and RWC sometimes also exhibited substantially non-linear behaviour over the range studied. In the five species studied, the slope was smaller than unity by a factor of a few, indicating a steeper decrease in H_c_^CR^ than in RWC. Non-linearity was possibly caused by changes in leaf structure, porosity and and/or leaf area during dehydration. At the levels of mechanical stress we worked with, compressed leaves likely dehydrate under a no-slip condition (with the epidermis firmly adhered to the compression plate), so the compressed region probably does not shrink in area during dehydration. In principle, decreases in RWC with no change in H_c_^CR^, as observed in *Fraxinus griffithii* (Fig. 3 D2), could result from leaf area shrinkage outside the squeezed region. Additionally, the steeper decline in H_c_^CR^ than in RWC suggests a decrease in leaf porosity during dehydration. Possibly, the latter is due to an increased ability of the cell walls to bend with decreasing turgor, which leads to progressive filling of the intercellular space during dehydration. Thus, while H_c_^CR^ may be correlated with leaf water content, care must be taken before assuming the linearity of this relation and the ranges over which it may be valid.

Creep and SR_SL_ experiments revealed non-linear relations between h_c_^CR^ and −σ^SRSL^ with variations in the area and shape of the hysteresis loop (Figs. 4 & 5). While we did not investigate the source of this behaviour, it is possible that these hydration hysteresis loops are caused by capillary effects and/or hysteresis in the stress-strain relations of cell wall materials and geometry. In the case of material hysteresis, energy may be dissipated by friction between cellulose microfibrils in a mechanism analogous to the deformation of rubbers (Joule, 1859). Capillary sorption hysteresis may play a role in the hydration of cell wall interstices, such as those described for wood (Barkas, 1942, Fredriksson and Thybring, 2019) and, more generally, for porous materials (Albers, 2014). Complex interactions between elastic and capillary hysteresis may also take place, as in high molecular weight polymers (Urquhart, 1929, Smith, 1947). Further research is needed to interpret the significance and mechanisms behind leaf hydration hysteresis loops.

The relation between h_c_^CR^ and −σ^SRSL^ and their hysteresis loop are relevant to the methods used to measure plant water transport. Commonly, plant ecophysiologists have studied plant hydration dynamics using invasive techniques that yield noisy data, carrying large uncertainty in the estimations of tissue conductance (Sack et al., 2002, Brodribb and Holbrook, 2003). Two main limitations of such methods are the use of excised tissues (which introduces inter-leaf variability and leads to potential artifacts in the measurements) and the lack of continuous measurements of leaf water potential. The last issue stems from the available techniques used to measure water potential: (i) the pressure bomb cannot be used quickly enough to track tissue rehydration with high temporal resolution; and (ii) thermocouple psychrometry has equilibration times which can be longer than the rehydration process. Recently, Bourbia et al. (2021) have addressed the issue by continuously estimating water potential from optical measurements of leaf petiole width; however, this method assumes a unvarying relation between petiole dimensions and water potential. In the case of uniaxially compressed leaves, we showed that this relation is most often not linear and is highly dependent on the direction of the hydration process. The lack of hysteresis during dehydration-rehydration cycles is sometimes assumed to estimate leaf hydraulic conductance (Brodribb and Holbrook, 2003), but previous research (Kamiya et al., 1963) and our present work indicate that hysteresis is present in dehydration-rehydration cycles of plant tissue. Thus, our results highlight long-standing issues and assumptions in the study of plant hydration dynamics and suggest an accessible method to approach them.

### In vivo measurements

A trial on a single potted *Avicennia marina* plant indicated that the technique is suitable to monitor changes in plant water status *in vivo*. In general, these results conformed to expectations of plant water status under changing light, and suggest that leaf squeeze-flow rheometry may be a valuable tool to monitor leaf thickness and turgor pressure non-invasively. Overall, the technique provides improved precision and temporal resolution when compared to the pressure chamber. However, temperature effects on the instrument dimensions currently limit the applicability of leaf squeeze-flow rheometry under field settings. Future efforts may develop the technique to be suitable for field conditions.

## Conclusion

Leaf squeeze-flow rheometry was applied as a technique to study plant hydration dynamics. The measurement paradigms of creep and stress relaxation were found to be useful means of monitoring changes in leaf relative water content and turgor pressure during dehydration-rehydration cycles, establishing a simple method to track leaf water status non-invasively. Our results and an idealised average-cell model of leaf uniaxial compression suggest that the leaf stiffness during compression is strongly dependent on the tissue osmotic pressure. While here we focused on static uniaxial compression under hydrostatic equilibrium, the study of non-equilibrium states during compression, as well as dynamic mechanical analyses such as oscillatory sweep tests, may provide further insights into plant water movement and broaden the applications of leaf squeeze-flow rheometry. Our findings may stimulate the development of more sophisticated actuated plant sensors with potential applications in productive and natural systems.

## Acknowledgements

We thank John Passioura for experimental support and advice and for his comments on the manuscript; Jacques Dumais for a discussion on the effects of leaf uniaxial compression on cell water status; Javier Merrill for comments on the manuscript; and Aria Carroll for her assistance in the experiments. TIF was funded by the Becas Chile program by ANID, Scholarship 72180256. This research was funded using TIF’s personal funds and supported by the Australian Research Council Discovery Project, Grant DP180102969.

## Author contributions

TIF conceived the hypotheses and designed the instrument. TIF designed and planned the experiments with suggestions from OB, CJB and MCB. TIF performed the experiments. TIF analysed the data and wrote the manuscript with suggestions from JW, OB, CJB and MCB.

## Footnotes

1 The pressure required to stop water diffusion through a semipermeable membrane separating a solution from pure water, equal to minus one times the osmotic potential. We use osmotic pressure throughout.

2 In preliminary experiments, we determined this pressure was enough to flatten the leaf lamina without causing visible damage to the tissue. In our system, this pressure is equivalent to a measuring force of 3.2 N; in comparison, the measuring force provided by the ratchet stop of the micrometer is 5-10 N.

3 In response to sudden compression, cells within porous tissues conserve volume by increasing width while reducing height. Assuming that the compressed sample is conserved in area because of friction against the pad, cells in non-porous tissues do not have the possibility of changing width, so their geometry is expected to change little during compression. The average cell would, however, conserve volume by increasing width while reducing height, as sketched.

## References

Albers, B. 2014. Modeling the hysteretic behavior of the capillary pressure in partially saturated porous media: a review. Acta Mechanica, 225, 2163–2189.

Barkas, W. W. 1942. Wood water relationships—VII. Swelling pressure and sorption hysteresis in gels. Transactions of the Faraday Society, 38, 194–209.

Beauzamy, L., Nakayama, N. & Boudaoud, A. 2014. Flowers under pressure: ins and outs of turgor regulation in development. Annals of Botany, 114, 1517–1533.

Bidhendi, A. J. & Geitmann, A. 2019. Methods to quantify primary plant cell wall mechanics. Journal of Experimental Botany, 70, 3615–3648.

Borsuk, A. M., Roddy, A. B., ThÉroux-Rancourt, G. & Brodersen, C. R. 2022. Structural organization of the spongy mesophyll. New Phytologist, n/a.

Bourbia, I., Pritzkow, C. & Brodribb, T. J. 2021. Herb and conifer roots show similar high sensitivity to water deficit. Plant Physiology, 186, 1908–1918.

Brodribb, T. J. & Holbrook, N. M. 2003. Stomatal closure during leaf dehydration, correlation with other leaf physiological traits. Plant Physiology, 132, 2166–2173.

De Belie, N., Hallett, I. C., Harker, F. R. & De Baerdemaeker, J. 2000. Influence of Ripening and Turgor on the Tensile Properties of Pears: A Microscopic Study of Cellular and Tissue Changes. Journal of the American Society for Horticultural Science jashs, 125, 350–356.

Engmann, J., Servais, C. & Burbidge, A. S. 2005. Squeeze flow theory and applications to rheometry: A review. Journal of Non-Newtonian Fluid Mechanics, 132, 1–27.

Franks, P. J., Cowan, I. R. & Farquhar, G. D. 1998. A study of stomatal mechanics using the cell pressure probe. Plant, Cell & Environment, 21, 94–100.

Fredriksson, M. & Thybring, E. E. 2019. On sorption hysteresis in wood: Separating hysteresis in cell wall water and capillary water in the full moisture range. PLOS ONE, 14, e0225111.

Geitmann, A. 2006. Experimental Approaches Used to Quantify Physical Parameters at Cellular and Subcellular Levels. American Journal of Botany, 93, 1380–1390.

Gibson, L. J., Ashby, M. F. & Easterling, K. E. 1988. Structure and mechanics of the Iris leaf. Journal of Materials Science, 23, 3041–3048.

Gibson, L. J., Ashby, M. F., Schajer, G. S. & Robertson, C. I. 1982. The mechanics of two-dimensional cellular materials. Proceedings of the Royal Society of London. A. Mathematical and Physical Sciences, 382, 25–42.

Green, P. B. 1962. Mechanism for Plant Cellular Morphogenesis. Science, 138, 1404–1405.

Green, P. B. & Stanton, F. W. 1967. Turgor Pressure: Direct Manometric Measurement in Single Cells of Nitella. Science, 155, 1675–1676.

Hamant, O., Heisler, M. G., JÖnsson, H., Krupinski, P., Uyttewaal, M., Bokov, P., Corson, F., Sahlin, P., Boudaoud, A., Meyerowitz, E. M., Couder, Y. & Traas, J. 2008. Developmental Patterning by Mechanical Signals in <em>Arabidopsis</em>. Science, 322, 1650–1655.

Hüsken, D., Steudle, E. & Zimmermann, U. 1978. Pressure Probe Technique for Measuring Water Relations of Cells in Higher Plants 1. Plant Physiology, 61, 158–163.

Jackman, R. L., Marangoni, A. G. & Stanley, D. W. 1992. THE EFFECTS OF TURGOR PRESSURE ON PUNCTURE AND VISCOELASTIC PROPERTIES OF TOMATO TISSUE. Journal of Texture Studies, 23, 491–505.

Joule, J. P. 1859. V. On some thermo-dynamic properties of solids. Philosophical Transactions of the Royal Society of London, 149, 91–131.

Kamiya, N., Tazawa, M. & Takata, T. 1963. The relation of turgor pressure to cell volume inNitella with special reference to mechanical properties of the cell wall. Protoplasma, 57, 501–521.

Lin, T.-T. & Pitt, R. E. 1986. RHEOLOGY OF APPLE AND POTATO TISSUE AS AFFECTED BY CELL TURGOR PRESSURE. Journal of Texture Studies, 17, 291–313.

Lockhart, J. A. 1965. An analysis of irreversible plant cell elongation. Journal of Theoretical Biology, 8, 264–275.

Peleg, M. 1979. CHARACTERIZATION OF THE STRESS RELAXATION CURVES OF SOLID FOODS. Journal of Food Science, 44, 277–281.

Philip, J. R. 1958. The Osmotic Cell, Solute Diffusibility, and the Plant Water Economy. Plant physiology, 33, 264–271.

Sack, L., Melcher, P. J., Zwieniecki, M. A. & Holbrook, N. M. 2002. The hydraulic conductance of the angiosperm leaf lamina: a comparison of three measurement methods. Journal of Experimental Botany, 53, 2177–2184.

Savitzky, A. & Golay, M. J. E. 1964. Smoothing and Differentiation of Data by Simplified Least Squares Procedures. Analytical Chemistry, 36, 1627–1639.

Scholander, P. F., Bradstreet, E. D., Hemmingsen, E. A. & Hammel, H. T. 1965. Sap Pressure in Vascular Plants. Negative hydrostatic pressure can be measured in plants, 148, 339–346.

Skotheim Jan, M. & Mahadevan, L. 2005. Physical Limits and Design Principles for Plant and Fungal Movements. Science, 308, 1308–1310.

Smith, S. E. 1947. The Sorption of Water Vapor by High Polymers. Journal of the American Chemical Society, 69, 646–651.

Tyree, M. T. & Hammel, H. T. 1972. The Measurement of the Turgor Pressure and the Water Relations of Plants by the Pressure-bomb Technique. Journal of Experimental Botany, 23, 267–282.

Urquhart, A. R. 1929. 15—THE MECHANISM OF THE ADSORPTION OF WATER BY COTTON. Journal of the Textile Institute Transactions, 20, T125–T132.

Weatherley, P. E. 1950. STUDIES IN THE WATER RELATIONS OF THE COTTON PLANT. New Phytologist, 49, 81–97.

Yang, D., Li, J., Ding, Y. & Tyree, M. T. 2017. Experimental evidence for negative turgor pressure in small leaf cells of Robinia pseudoacacia L versus large cells of Metasequoia glyptostroboides Hu et W.C. Cheng. 2. Hofler diagrams below the volume of zero turgor and the theoretical implication for pressure-volume curves of living cells. Plant, Cell & Environment, 40, 340–350.

