## Supplement for "Measurement of plant water status via static uniaxial compression of the leaf lamina"

**Supplementary material**

**Table S1.** Species and sample sizes used for the experiments conducted in this study. Experiment numbers correspond as: (1) creep, SR_SL_ and SR_SS_ under constant water status; (2) dead leaf test; (3) creep and SR_SL_ under changing water status; (4) creep and SR_SL_ on a living plant.

| Species | Family | Order | Experiment | Sample size |
| --- | --- | --- | --- | --- |
| Populus nigra L. | Salicaceae | Malpighiales | 1 | 1 |
| Salvia officinalis L. | Lamiaceae | Lamiales | 1 | 1 |
| Quillaja saponaria Molina | Quillajaceae | Fabales | 2 | 3 |
| Arbutus unedo L. | Ericaceae | Ericales | 3 | 1 |
| Callistemon viminalis (Sol ex. Gaertn.) G. Don | Myrtaceae | Myrtales | 3 | 3 |
| Corymbia citriodora (Hook.) K. D. Hill & L. A. S. Johnson | Myrtaceae | Myrtales | 3 | 1 |
| Fraxinus griffithii C. B. Clarke | Oleaceae | Lamiales | 3 | 1 |
| Grevillea olivacea A. S. George | Proteaceae | Proteales | 3 | 1 |
| Ligustrum lucidum W. T. Aiton | Oleaceae | Lamiales | 3 | 3 |
| Podocarpus elatus R. Br. ex Endl. | Podocarpaceae | Pinales | 3 | 1 |
| Quercus ilex L. | Fagaceae | Fagales | 3 | 1 |
| Tristaniopsis laurina (Sm.) Peter G. Wilson & J. T. Waterh. | Myrtaceae | Myrtales | 3 | 3 |
| Avicennia marina subsp. australasica (Walp.) J. Everett | Acanthaceae | Lamiales | 4 | 1 |

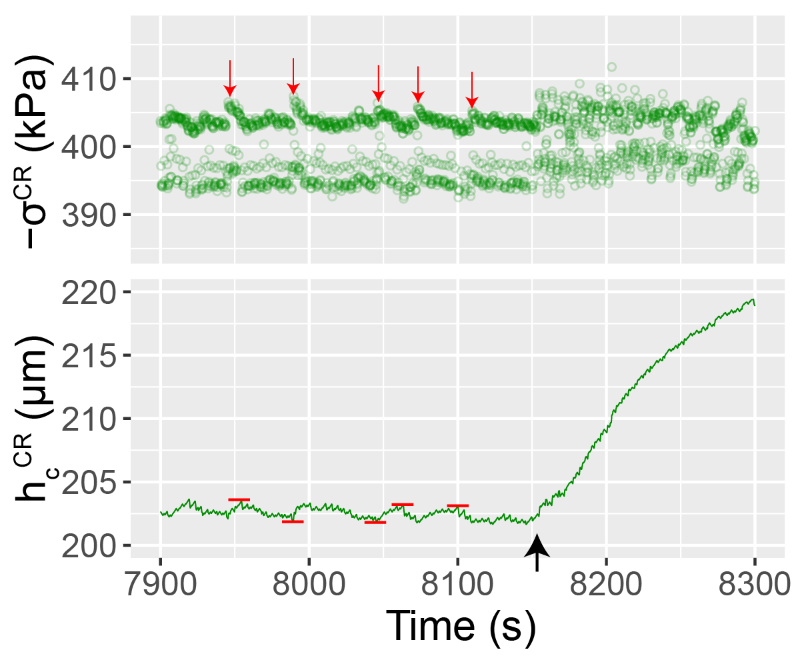

**Figure S1.** Example of the backlash of the instrument during a creep test conducted at 400 kPa. Red downward arrows indicate the approximate times at which the motor took an effective compressive step during dehydration, increasing the applied stress. Red horizontal bars denote the recorded thickness at the times indicated by the arrows. Backlash corresponds to the approximate vertical distance between the red horizontal bars, which in this system is *c*. 2 µm. The black upward arrow corresponds to the time at which the dehydrating sample was recut under water. Note the change in direction of the trend in pressure, and the absence of backlash during fast rehydration.
